# In vivo microvascular flow quantification in the mouse brain using Row-Column Ultrasound Localization Microscopy and directed graph analysis

**DOI:** 10.1101/2025.09.17.676732

**Authors:** Adrien Bertolo, Jeremy Ferrier, Oscar Demeulenaere, Alexandre Dizeux, Tanguy Delaporte, Bruno Osmanski, Mickael Tanter, Mathieu Pernot, Thomas Deffieux

**Author notes:** These authors contributed equally to this work (Mathieu Pernot, Thomas Deffieux).

## Abstract

Brain perfusion relies on a complex vascular network of arteries, veins, and capillaries to meet its constant demand for oxygen and nutrients. Disruption of this microvascular system is a hallmark of many neurological disorders, including small vessel disease, stroke, and brain tumors. As such, high-resolution in vivo imaging of cerebral microvascular flow and structure remains critical to understanding these pathologies. Among them, Ultrasound Localization Microscopy (ULM) allows noninvasive imaging of microvascular network at subwavelength resolution using injected microbubbles, but the approach remains mainly limited to 2D imaging with few volumetric implementations. In this study, we explore in vivo transcranial 3D ULM of the mouse brain using Row-Column Arrays (RCA) and introduce an analysis framework to build a flow-directed vascular graph from the ULM microbubble tracking data, allowing to differentiate between subgraphs of artery-like and vein-like vascular segments. Combined with Allen-based and Radius-based segmentations, we extract metrics including cerebral blood flow (CBF), microbubble (MB) velocity, flowrate, CBV fraction, vascular segment length and tortuosity in sixty-six different vascular classes. This high-sensitivity framework enables in vivo microvascular imaging and quantification in mice and provides a scalable platform for preclinical neurovascular studies in health and disease.

## Introduction

Preclinical neuroimaging has become an indispensable tool in neuroscience research for imaging the brain’s vascular network, a highly intricate system responsible for regulating cerebral blood flow, oxygen and nutrient delivery, and metabolic waste clearance. This network is fundamental to maintaining brain health, yet many aspects of its structure and function remain poorly characterized. Emerging evidence suggests that alterations in microvascular structure and function may serve as early biomarkers for a range of neurological disorders^1^. For instance, small-vessel diseases and neurodegenerative disorders - such as Alzheimer’s disease - are often preceded by microvascular alterations^2^, and such alterations may contribute to disease progression before overt symptoms appear. In vascular stroke, phenomena such as vasospasms, aneurysms, and delayed ischemia disrupt blood flow across multiple spatial and temporal scales and are critical areas of study for developing and testing effective recovery and intervention strategies^3^. In brain tumors, the formation of aberrant and complex vascular networks is essential for supporting rapid tumor growth and survival. Imaging these networks is not only vital for understanding tumor biology but also for developing targeted therapeutic approaches to disrupt their blood supply and limit progression.

Several imaging modalities have been developed for neurovascular mapping, including magnetic resonance angiography (MRA)^4,5^ and computed tomography angiography (CT) using contrast agents but also optical imaging techniques^6^. However, MRA and CT imaging techniques are limited in their ability to provide both high-resolution imaging of microvascular architecture and blood flow dynamics, while optical imaging is restricted to surface and cannot image deeper brain regions. While advanced ex vivo techniques, such as optical clearing methods (e.g., iDISCO, CLARITY)7 and synchrotron-based phase-contrast tomography^8–10^, excel in providing unparalleled spatial resolution for microvascular imaging, they come with significant limitations. These techniques are highly effective for detailed structural visualization of cerebral vasculature, often achieving resolutions at the sub-micron to micron scale but are inherently restricted to fixed tissue samples and thus lack the ability to capture dynamic flow information, such as blood velocity or cerebral blood flow (CBF). These ex-vivo approaches are not suitable for longitudinal studies, as they require tissue extraction and processing, preventing repeated measurements in alive subjects.

These limitations highlight the need for novel imaging modalities that can provide both high-resolution structural imaging and dynamic functional assessment of the brain’s microvasculature in vivo. Ultrasound Localization Microscopy (ULM) is an emerging imaging technique that can map microvascular flow at high spatial and temporal resolution. ULM relies on the imaging, localization, and tracking of FDA-approved gas microbubbles circulating in the blood flow^11^ using ultrafast ultrasound imaging. The technique has been applied to map cerebral microvascular flow in rodents and in humans using 2D imaging approaches allowing subwavelength resolution in depth and providing quantitative velocity estimates^12^. Recently, functional ULM (fULM) has been introduced to map functional hyperemia in the rodent brain at high spatial resolution and in time^13^, access to high spatial resolution maps allows to probe different vascular compartments and further investigate the neurovascular coupling along different scales in depth.

While Ultrasound Localization Microscopy (ULM) has significant potential to increase our understanding of cerebral microvasculature, its generic two-dimensional implementation presents important limitations. Single-plane imaging not only limits the spatial regions that can be examined simultaneously but also introduces systematic biases in velocity measurements, as only two components of the three-dimensional velocity vector can be reliably captured. These constraints have motivated the development of three-dimensional (3D) ULM approaches using dense ultrasonic matrix arrays, composed of thousands of piezoelectric elements arranged in two-dimensional configurations instead of linear configuration for rodent brain imaging including stroke and tumor models ^14–18^. Despite these early proofs of concepts, matrix arrays still face substantial practical and technical challenges. Manufacturing these arrays remains costly and technically demanding, particularly at the high frequencies above 10 MHz typically required for preclinical neuroimaging applications such as rodents, and a fundamental trade-off exists between acoustic element pitch, which governs sidelobe levels and overall sensitivity, and the practical challenges of array fabrication and precision^19^. This compromise makes scaling to larger fields of view particularly challenging without exponentially increasing the number of elements, thereby escalating costs and system complexity. While electronic multiplexers can mitigate some of these requirements by dynamically switching element^17,20^, this approach inevitably reduces the effective volume rate due to circuit switching times and the sequential addressing of array elements, compromising microbubble tracking efficiency^21^ in the case of ULM and overall image quality^22^.

To address these limitations, Row-Column Array (RCA) probes have emerged as a promising alternative for three-dimensional high volume rate ultrasound imaging. Conceptually, RCA probes consist of two overlapped orthogonal one-dimensional arrays which offers a simplified, yet efficient approach compared to matrix arrays for 3D ultrasound imaging. RCA enables larger field-of-view imaging with reduced complexity while maintaining improved sensitivity and can be manufactured with finer pitches even at high frequencies. RCA enables a significant reduction of channels compared to matrix arrays (N+N rather than N^2^, N being the number of elements along one side of the array), which remains compatible with standard 256-channel preclinical ultrasound scanners. The versatility of RCA probes in preclinical neuroimaging has been demonstrated across various modalities, including Doppler imaging^23^, functional ultrasound imaging (fusi)^24^ and non-linear contrast imaging of gas vesicles^25^. Their potential has been explored in proof-of-concept studies of 3D ultrasound localization microscopy (ULM)^26,27^ including recently 3D imaging of the mouse brain by Wu and colleagues^28^as and the rat brain by Sun et al^29^. While RC-ULM shows clear promise for three-dimensional ULM imaging in rodent brains, further advancements in acquisition protocols, reconstruction methods, and analytical frameworks are still needed toward achieving a high-throughput, benchside microvascular flow quantification technique suitable for in vivo preclinical studies.

Our study explores transcranial 3D Ultrasound Localization Microscopy (ULM) in mouse models using a dedicated Row-Column Array (RCA) probe through the combination of optimized acquisition sequences, advanced reconstruction algorithms, and automated flow-directed graph-based analysis of vessels. We first demonstrate the capability of this approach to reconstruct the 3D cerebral microvasculature through the skull with high sensitivity and spatial resolution, including the local estimation of blood velocities, the 3D vessel reconstruction and its registration to the Allen Brain Atlas. We further show that these data can enable the construction of a flow-directed vascular graph which locally encodes flow direction and can then be used for labeling artery-like and vein-like vascular segments based on local branching and flow gradient parameters. By combining directed graph with local vessel radii estimation, we further extend the analysis to the automated quantification of dynamic features (cerebral blood flow, microbubble velocity) and vessel structural characteristics (segment length, tortuosity) across both distinct vascular compartments and anatomical regions across the brain.

## Material and Methods

### Ethics

The institutional and regional committees approved the study for animal care (Comité d’éthique pour l’expérimentation animale \#59 - “Paris Centre et Sud,” Protocole \# 2017-23) and adhered to the ARRIVE recommendations. All animals received humane care in accordance with the European Union Directive of 2010 (2010/63/EU).

Mice were naive, seven weeks old at the time of the experiment, and randomly placed in the cage; none were excluded. The individual animal is the experimental unit in this investigation.

### Animal preparation

Five male C57Bl/6 mice (Janvier Labs; Le Genest St Isle, France) were utilized in this study. The mice were housed under controlled conditions (22 1 ° C, 60% relative humidity, 12/12h light/dark cycle, ad libitum food and drink) for a week prior to the start of the experiment. The mice were anesthetized with an initial administration of ketamine (Imalgene, 100 mg/kg) and Xylazine (Rompun) intramuscularly to render them unconscious, and anesthesia was maintained with a 1.5% isoflurane supply and the animal physiology was monitored. The animals were then placed in a stereotaxic frame. The skin and the skull were kept intact. The eyes of the mouse were protected using an ointment (Ocry-gel, TVM, UK). Body temperature was controlled with a rectal probe connected to a heating pad set at 37°C. Respiration and heart rate were monitored using a PowerLab data acquisition system with the LabChart software (ADInstruments, USA). Additional IP ketamine/xylazine doses (25 mg/kg and 2.5 mg/kg, respectively) were infused intermittently (every 90 to 120 min), as deemed necessary based on changes in physiological parameters. ULM was performed using Sonovue bolus injections in the tail vein.

### RC-ULM acquisition

#### Row-Column Array, sequences and scanner

A piezoelectric 15 MHz row-column array (Iconeus, Paris, France) consisting of 80 rows and 80 columns with a 0.110-millimeter pitch was used to perform the 3D power Doppler imaging and 3D Ultrasound Localization Microscopy acquisitions. The array was connected to a 256-channel functional ultrasound scanner (Iconeus One, Iconeus, Paris, France) driving the probe at 12.5 MHz instead of 15 MHz to limit the ultrasonic wave aberrations induced during the propagation through the skull bone.

For ultrafast power Doppler, the imaging sequence was defined by transmitting 20 plane waves along the rows followed by 20 plane waves with the columns (angular range of 15°). A four-half-cycle pulse with a duty cycle of 67% and a tension of 25 V was used with a volume rate of 500 Hz. Each block of 200 images (0.4 s) was acquired at 1 Hz to limit the probe heating.

The probe was positioned perpendicular to the animal head’s surface, and a four-axis motor module (Iconeus, Paris, France) was used to ensure proper positioning of the probe using a real-time 3-view Doppler imaging mode.

The ultrafast RC-ULM sequence was defined by transmitting 14 plane waves along the rows followed by 14 plane waves along the columns at a PRF of 28 kHz, yielding a high volume rate of 1000Hz (angular range, 10°). A two-half-cycle pulse was sent at a frequency of 12.5 MHz and a tension of 20 V. The ultrafast sequence comprised 500 volumetric images acquired in 500 milliseconds, repeated 400 times for a total accumulation time of 200 seconds. The acquisition time was 6 minutes and 40 seconds, with 0.5-second intervals between each block of 500 ultrafast volumes.

#### Beamforming and XDoppler reconstruction

RCA beamforming was implemented on two GPUs (A6000 boards, Nvidia, USA) on the scanner and optimized to reach real-time 3D beamforming of the ultrafast volumes including GPU-based clutter filtering. This allowed to perform both the live power Doppler volumetric imaging with a 3-view rendering for the positioning but also perform the online reconstruction of the microbubble volumes using the XDoppler scheme^30^ down to the prelocalization of microbubbles.

More specifically, the volume RC (transmission along Rows and reception along Columns) was beamformed using a conventional Delay-And-Sum beamformer optimized for the dual GPUs. The beamformer grid was set to (λ×λ×λ/2) and the different angles transmitted along the row were coherently compounded. A SVD clutter filter was then applied on this stack with an ensemble length of 500 frames. The same processing was applied to the CR volume (transmission along Columns and reception along Rows). The filtered RC and CR stacks were then cross correlated with a window size of 5 frames for ensemble length using the XDoppler scheme. This correlation approach allows to reduce the signal from the lobes, compared to a synthetic summation approach like orthogonal plane wave compounding (OPW)^30^.

Within each ultrafast block, MB were prelocalized in real time using a local maximum algorithm and a neighborhood of (5×5×5) voxels around each maximum was extracted and stored on disk for further processing and refined localization. All those steps were implemented on the GPU in Cuda and running in real time.

### 3D ULM processing

The localizations were then refined offline using a 3D interpolation with a gaussian kernel of each neighborhood volumes followed by a 3D gaussian filter. The sub voxel localization was performed using a 3D paraboloid fit, the resulting position table was stored and a tracking algorithm was used to pair microbubbles from one frame to another within given physical constraints on pairing distance and velocity ranges using a Hungarian linker^31^. The tracks were subsequently rasterized to generate microbubble (MB) count volumes and MB velocity component volumes (10 μm isotropic).

For visual distinction between cortical arterioles and venules, we also reconstructed a signed MB velocity map in which we flipped the sign of velocity norm for every voxel whose axial velocity was negative.

### Comparison with Multiplexed Fully Populated Matrix arrays (MUX-FPM)

We compared the RC-ULM scheme with a MUX-FPM, centered at 15 MHz, on the same animal. After the RC-ULM acquisition, a pause of 30 min allowed the mouse to recover and to clear the remaining MB from the circulation. During this time, the RCA probe was replaced with the MUX-FPM array. A second imaging session of the same acquisition duration (240 blocks of 0.5 each) was then performed using the same ultrasound frequency and the same voltage. The imaging sequence was adapted from the work of Chavignon and colleagues ^20^ where 10 transmit/receive combinations were repeated per tilted plane wave. The PRF was fixed at 12 kHz and 6 tilted plane waves were transmitted using the following angular sequence: (-5°, 0°); (0°; 0°); (0°, 5°); (0°, -5°); (0°, 0°); (0°, 5°). The resulting framerate of 200 Hz allowed the accumulation of 100 compounded frames per 0.5s block. The SVD cutoff for the clutter filter was set to 10/100 after optimization. After localization and tracking, we generated volumetric density and velocity maps (resolution 10 µm). The resulting ULM volume was realigned to the RCA volume automatically using the transformation found by intensity based registration between the two corresponding power Doppler volumes. The MEX-FPM ULM density map was resampled in the RCA space for side-by-side visual comparisons (see supplementary figure 6).

### Spatial resolution assessment

To evaluate the spatial resolution of the proposed vascular imaging modality, we first measured the Fourier Shell Correlation (FSC) between two MB count volumes, obtained by rasterizing two independent sub-groups of MB tracks split from the same acquisition. Another relevant assessment of the spatial resolution was then performed by evaluating the capacity to separate the MB velocity profiles from small and close cortical vessels in the mouse brain. The velocity profile selection was performed in a region of interest containing two penetrating arterioles using Amira software. Velocity profiles were extracted on three different axial slices (thickness 20 µm), taken at three different depths to intersect different sub-branches of the same penetrating arteriole. Across each profile, the MB velocity was extracted using the Line Probe function of Amira Avizo, using a sampling step of 5 microns.

### Microvascular graph reconstruction and analysis

#### Graph construction from ULM maps

As a first step, the MB density volumes were filtered using a hessian-like 3D spatial filter to enhance vessels (3D Jerman enhancement filter^32^). The resulting volume was then thresholded and skeletonized to extract the vessel centerline coordinates. Radius estimation was performed on these centerline coordinates using the VesselVio^33^ toolbox. VesselVio allowed to build a graph structure representing the vascular tree, described as a list of edges (describing a vascular segment) connecting vertices with a connectivity of 2 between branching nodes (vertices with a connectivity of at least 3). Each edge was hence characterized by a list of coordinates associated to these vertices. Vascular segment morphological features such as radius, length, surface, volume and tortuosity features such as MB density and 3D MB velocity vector were extracted and stored in the graph data. The radius and centerline MB velocity estimations were used to further estimate the flowrate in each segment of the vascular graph, as proposed in our previous work^34^ which was also stored for each vascular segment in the graph data.

#### Graph orientation from local blood flow direction

To orient the graph of vascular segments, we took advantage of the locally measured flow direction and the graph geometry. The orientation of each edge followed the direction of the flow, meaning that the sign of the scalar product between the average MB velocity vector field and the orientation vector of each segment was always positive. By running the Breadth First Search (BFS) algorithm as implemented in the *igraph* library in Python on this directed graph object, perfused or drained territories could be easily revealed from any seed segments. A graph traversal algorithm was also implemented in order to reveal the tree structure of any cluster of connected components and store the branching level of each segment in this tree.

#### Artery-like and vein-like labeling from graph analysis

Then, vascular segments were classified in either artery-like or vein-like groups using a multistep multiscale approach. The first step, more global, consisted in identifying seed segments with a flowrate higher than 3 µL/min, supposed to be potential upstream initiators of perfused territories or downstream attractors of drained territories. We could then trace the path of blood flow circulation and identify the descendants (perfused segments) or ancestors (feeding segments) of each seed segment and estimate the flowrate gradient along the vascular path. Segments associated with a negative flowrate gradient along the vascular path were labelled as artery-like segments, whereas segments associated with a positive flowrate gradient were labelled as vein-like segments. The second step, more local, allowed us to confirm and to complement the results of the first step, and consisted in estimating for each node the difference between its outward connectivity (number of descendants) and its inward connectivity (number of ancestors). The nodes with a diverging behavior were detected with a positive branching difference, and their connected vascular segments were classified as artery-like. Alternatively, the nodes with a merging behavior detected with a negative branching difference, and their connected vascular segments classified as vein-like.

#### Vascular compartments labeling from vascular type and local diameters

To further classify the different parts of the directed vascular tree, we separated the different branches based on their diameters in three classes: large, medium and small segments. Diameter thresholds (∅_T1, ∅_T2) were chosen to obtain a clear distinction of the three classes of segments (S class: (∅<∅_T1) - M class: (∅_T1<∅<∅_T2) – L class: (∅>∅_T2), with ∅_T1 = 50 μm and ∅_T2 = 70 μm. These thresholds were chosen to group together penetrating arterioles and venules in the S class, arteries and veins including pial vessels in the M class, and major arteries and veins such as arteries from the polygon of Willis or veins from the different sinuses in the L class.

#### Template registration and Allen-based region labeling

The geometric transformation aligning the brain vascular volume to the Allen mouse brain atlas^35^ was automatically estimated using the Iconeus Brain Positioning System, as described by Nouhoum et al.^36^, and implemented in the IcoStudio software (Iconeus, Paris, France). This process involved aligning a power Doppler volume - extracted from the first 20 seconds of the continuous ULM acquisition - to a mouse brain power Doppler angiography template which was already pre-aligned with the Allen Brain atlas^35^. Once the optimal affine transformation was identified, it could be used for ULM quantification in any anatomical region from the Allen atlas.

Vascular segments were then assigned to different anatomical regions of interest based on the template registration results. We chose 11 regions in the study (see figure 6.i.). For each chosen region, we tested if the coordinates of the graph centerline points were inside or outside the closed mesh of said region using *intriangulation* ^37^. This further allowed us to classify the different vascular segments by region in addition to their diameter and vascular type.

#### Vascular metrics extraction by class

Vascular segments were classified according to three previously defined criteria: vascular type (artery-like or vein-like), anatomical regions (based on the 11 main Allen atlas areas), and vessel diameter (small, medium, or large), resulting in sixty-six distinct classes. For each class, we extracted in vivo metrics encompassing both morphology- and flow-related parameters. The morphology-related metrics^38^ included segment length, segment tortuosity, and the cerebral blood volume (CBV) fraction, while the flow-related metrics comprised microbubble (MB) velocity, flow rate, and cerebral blood flow (CBF).

The CBV fraction in each anatomical region was defined as the ratio of vascular volume to the total volume of the corresponding region. CBF was defined as the total flowrate delivered to the tissue per unit volume within an anatomical region of interest (µL/min/mm^3^).

### Data visualization

The 3D renderings of the MB density and signed velocity volumes (Figure 3 and Figure 4) were performed using the Amira software (Amira v.6.0.1 software, Visualization Sciences Group, USA). The 2D slices represented in the Figure 3 were reconstructed with a prototype of IcoStudio software featuring 3D ULM tracks visualization (Iconeus, Paris, France). The 3D vascular graph renderings were rendered using PyVista visualization toolkit (Figure 5) or using Matlab (Figure 6). Flowrate for artery-like subgraph was rendered in red whereas it was rendered in blue for vein-like subgraphs (Figure 5 and Figure 6).

**Figure 1:**
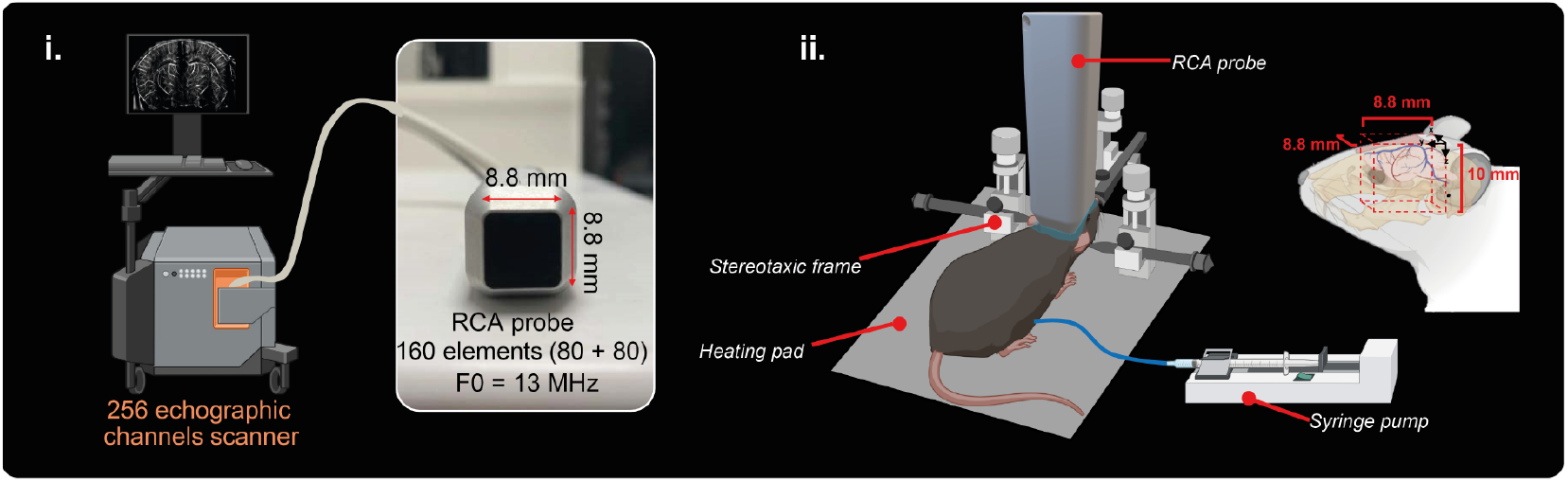
3D ULM acquisition setup with the RCA probe. i. Iconeus One driving the 160 elements RCA probe centered at 13 MHz. ii. Experimental setup for the 3D ULM experiment in the anesthetized mouse brain, with intact skull and skin.

**Figure 2:**
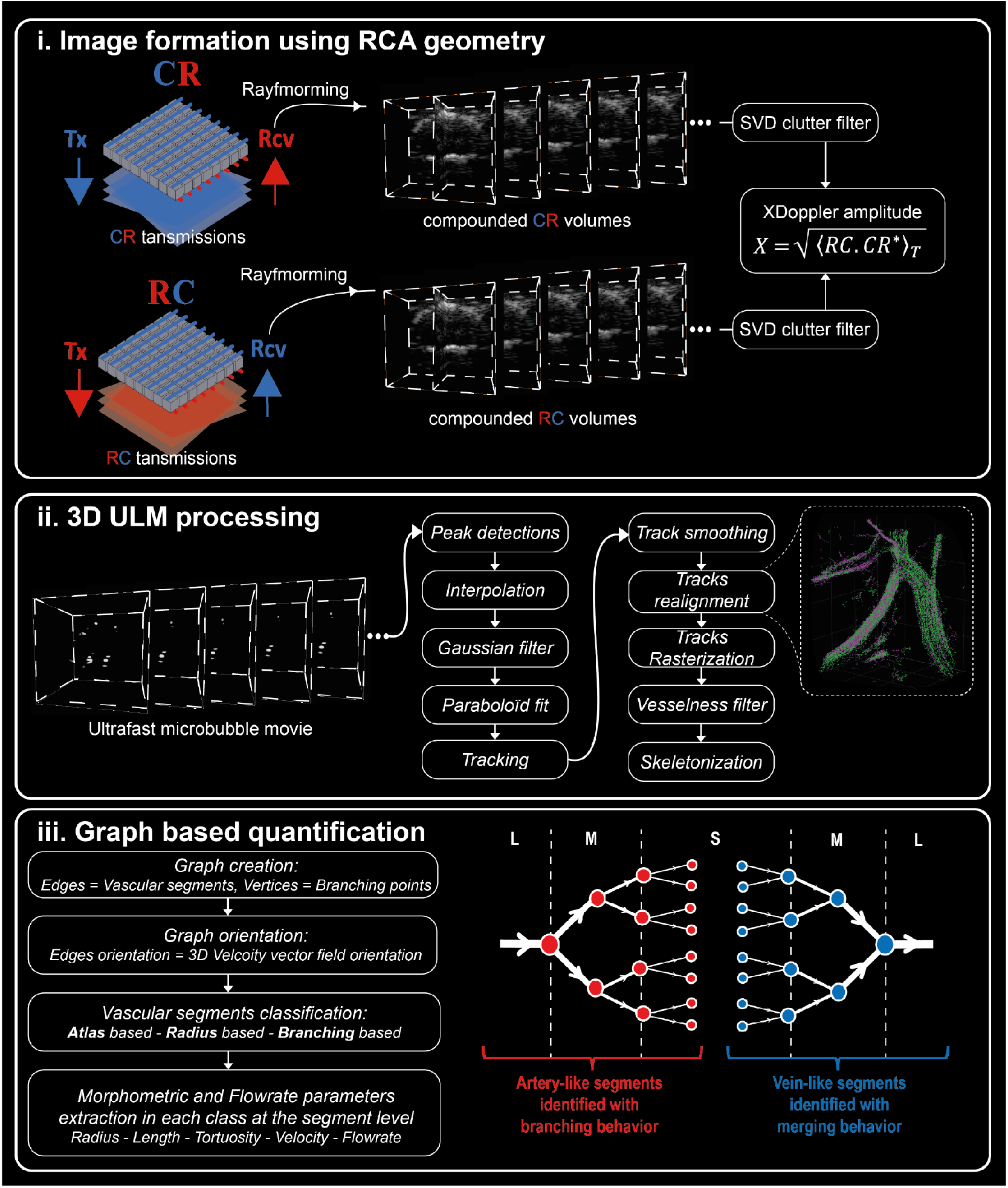
i. Synthetic focusing with the RCA probe geometry. The columns (C) are represented in blue, and the rows (R) are represented in red. To obtain an isotropic PSF, the final volume is reconstructed by combining the backscattered echoes acquired with the rows (after plane wave transmission through the columns) and the columns (after plane wave transmission through the rows) using the XDoppler strategy. ii. The ultrafast BMode movie of the MB is processed by the ULM processing pipeline. After tracking, the MB positions are used to correct the potential spatial drifts before rasterization. A vesselness filter is then applied on the resulting rasterized volume, before skeletonization. iii. Finally, a graph-based processing pipeline is applied to extract morphometric and flowrate parameters at the segment level in different classes of vascular segments. The first level of classification is defined by the anatomical segmentation, the second level of classification is based on diameter estimations: S class: (∅<∅_T1) - M class: (∅_T1<∅<∅_T2) – L class: (∅>∅_T2), and the last level of classification is based on the branching (artery-like segments) or merging (vein-like segments) behaviors observed on the directed graph.

**Figure 3:**
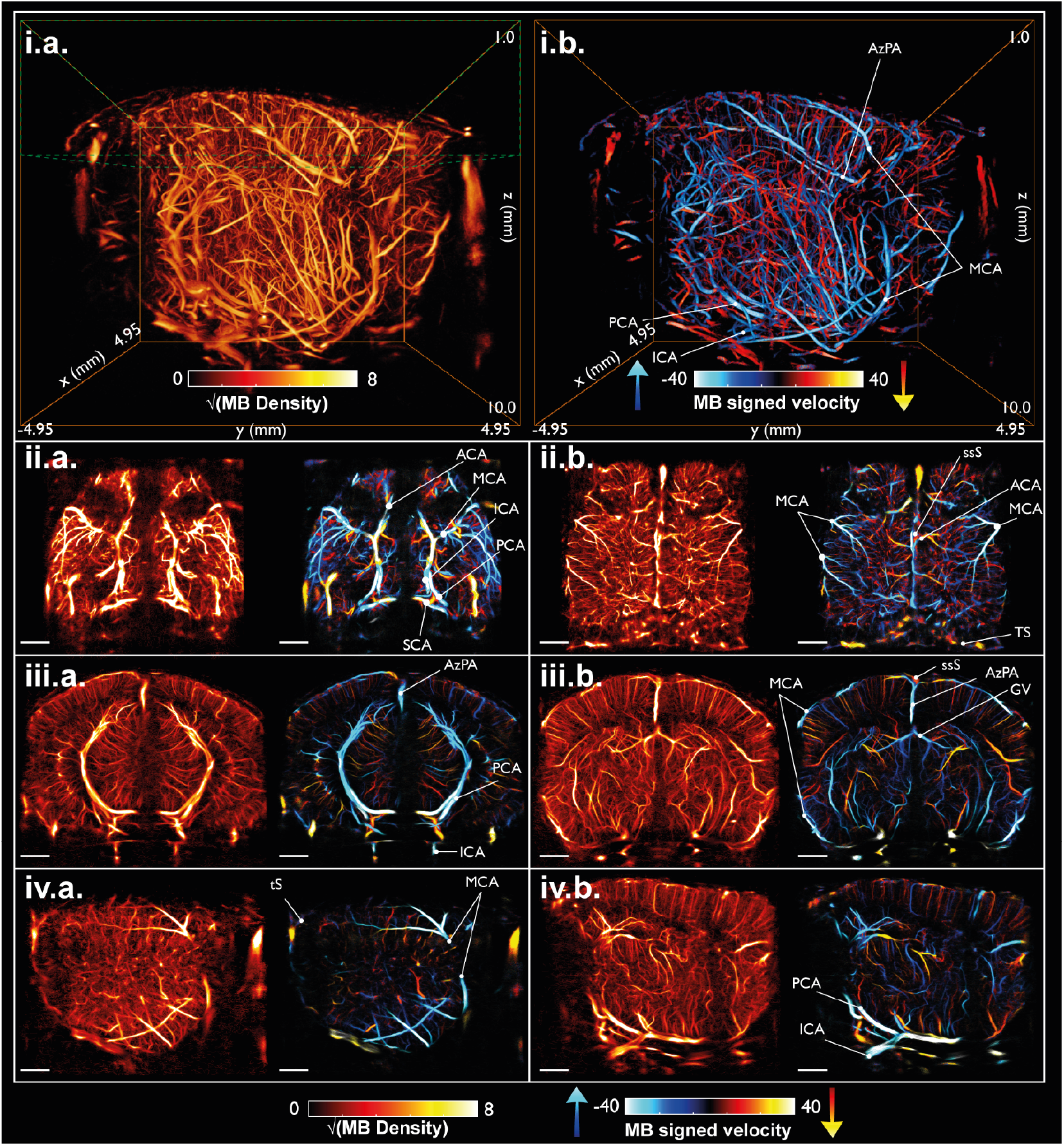
3D ULM visualization i. Sagittal Maximum Intensity Projection (MIP) of the 3D MB density volume (i.) and of the 3D signed MB velocity volume (ii.) are rendered with Amira software. The ULM tracks are rendered in the following 2D slices with IcoStudio software. The axial slices displayed in ii.a. and in ii.b. have a thickness of 2 mm. The scalebars represent 1 mm. The coronal and sagittal slices presented in iii. and in iv. have a slice thickness of 1.5 mm. Abbreviations: CoW : Circle of Willis ; ICA : Internal Cerbral Artery ; PCA : Posterior Cerebral Artery ; MCA : Medial Cerebral Artery ; Azc : Asygos ; SS : Sagittal Sinus ; TS: Transversal Sinus ; PSV : Posterior Superficial Vein ; GV : Galeno Vein.

**Figure 4:**
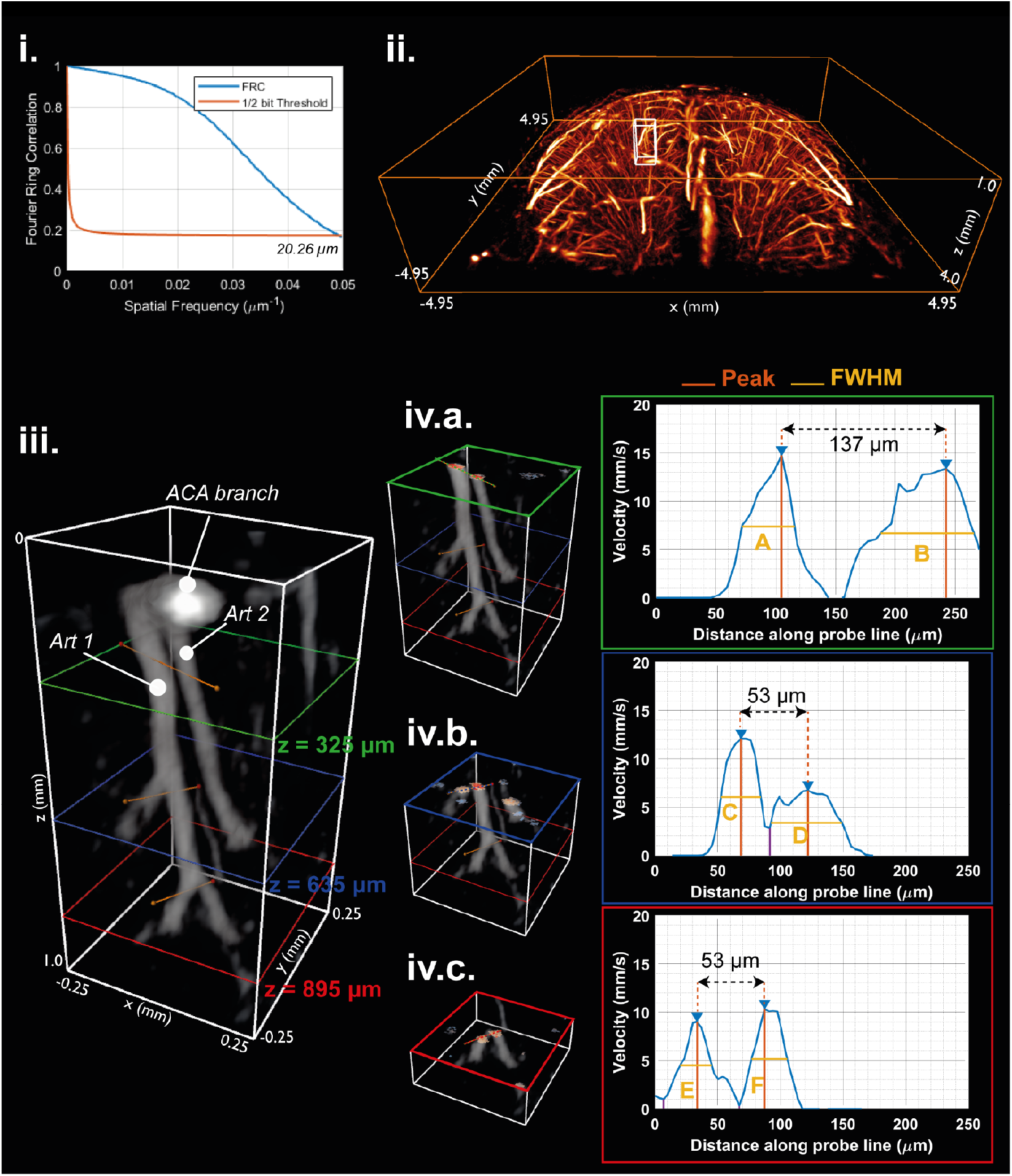
Spatial resolution assessment: Characterization of the MB velocity on different branching levels of the same penetrating arteriole. (i) The spatial resolution measured by the Fourier Shell Correlation is 20,26 µm (ii) Zoom of the MB density volume in the cortex of the mouse brain (corresponding to the green bounding box in Figure 3.i.). The penetrating arterioles ROI (white box) of size (4λ x 4λ x 10λ) was selected within the ACA territory. (iii) The selected region contains two arterioles (ArtA and ArtB). Three planes were placed at z = 325 µm (green), z = 635 µm (red) and z = 895 µm (blue). On each plane, the velocity intensity was extracted along the 1D profiles represented with the orange lines (ii). For each depth, a cropped view showing the selected plane is represented in (iii), and the velocity profile is shown in a transparent overlay. The corresponding velocity profile is plotted next to each cropped volume (iv). At z = 325 µm, the profiles crosses ArtA and ArtB and reveals a distance of 137 µm between them. At z = 635 µm and z = 895 µm, the velocity profile crosses new branches of ArtA, which are separated from 53 µm. The FWHM values are: A = 43.5 µm; B = 99 µm; C = 31.6 µm; D = 65 µm ; E = 25 µm; F = 29.1 µm

**Figure 5:**
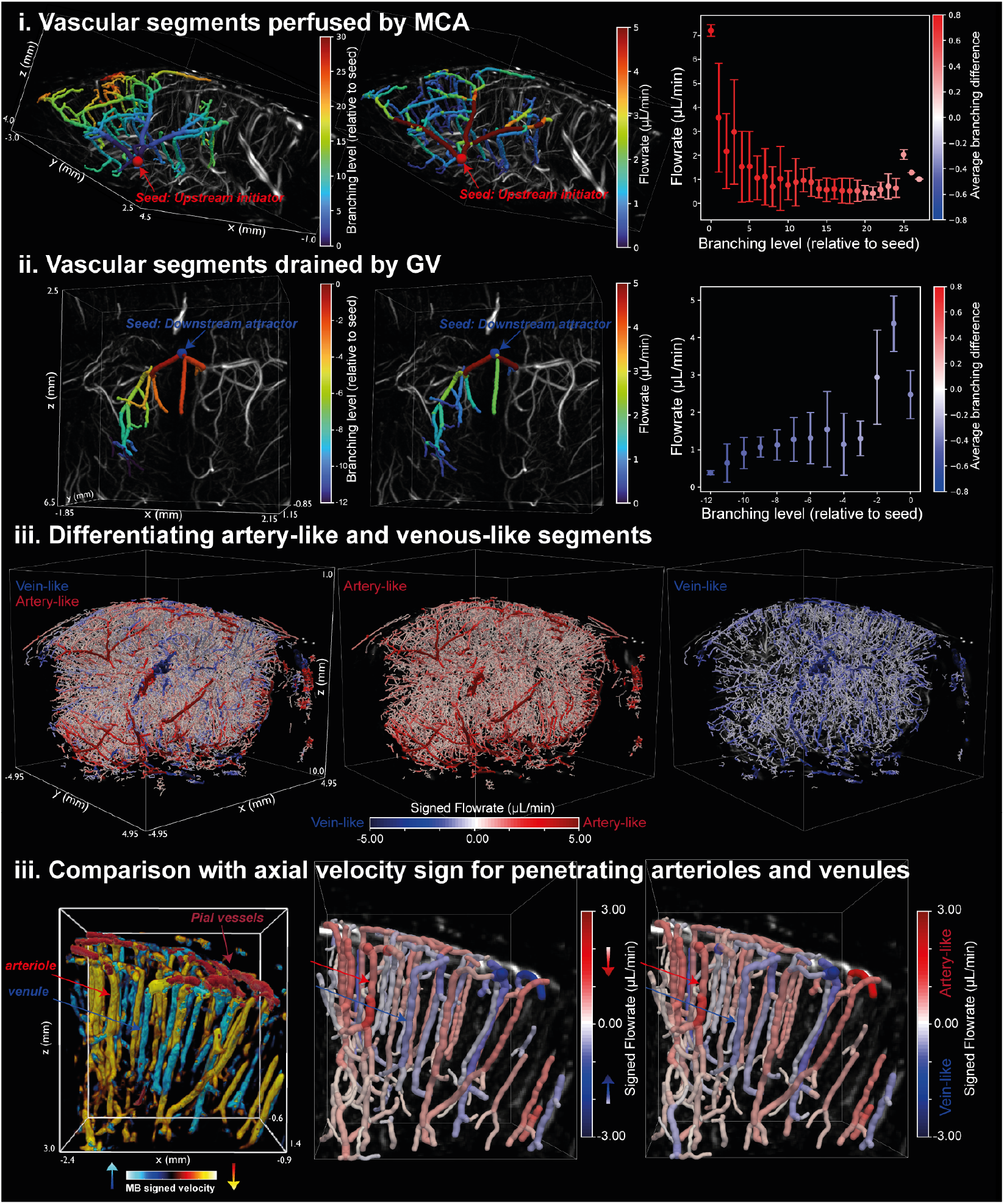
Mapping perfused and drained vascular territories, and labeling artery-like or vein like vascular segments using directed graph analysis. (i) The vascular territory perfused by the MCA is identified by selecting a seed segment (red sphere), upstream of the MCA bifurcation, and extracting all downstream (descendants) segments. Segments are represented over the MB density volume. Tube colors encode branching level (relative to the seed) on the left, and flowrate on the middle. The right plot shows a negative slope of average flowrate across branching levels, alongside a positive branching difference at the node level (red color code), reflecting the diverging pattern of artery-like vascular territories. (ii) In contrast, the drained vascular territory of the GV is identified by selecting a seed near the GV confluence node of the GV and tracing upstream (ancestor) segments. Here, segments merge at successive level, with increasing flowrate along the vascular path. The positive gradient is accompanied by a negative branching difference at the node level (blue color code), highlighting the converging behavior of vein-like vascular territories. (iii) The sign of flowrate renderings in the whole mouse brain is based on the branching behavior. The artery-like subgraph (red) is overlayed to the vein-like sub-graph (blue) on the left rendering, and each subgraph is represented separately on the middle and right renderings. (iv) The left rendering represents the MIP of the signed MB velocity volume in a small region of interest with penetrating arterioles and venules, in which a good match is observed between the sign assignment based on the axial MB velocity (middle) and the sign assignment based on the branching classification (right).

**Figure 6:**
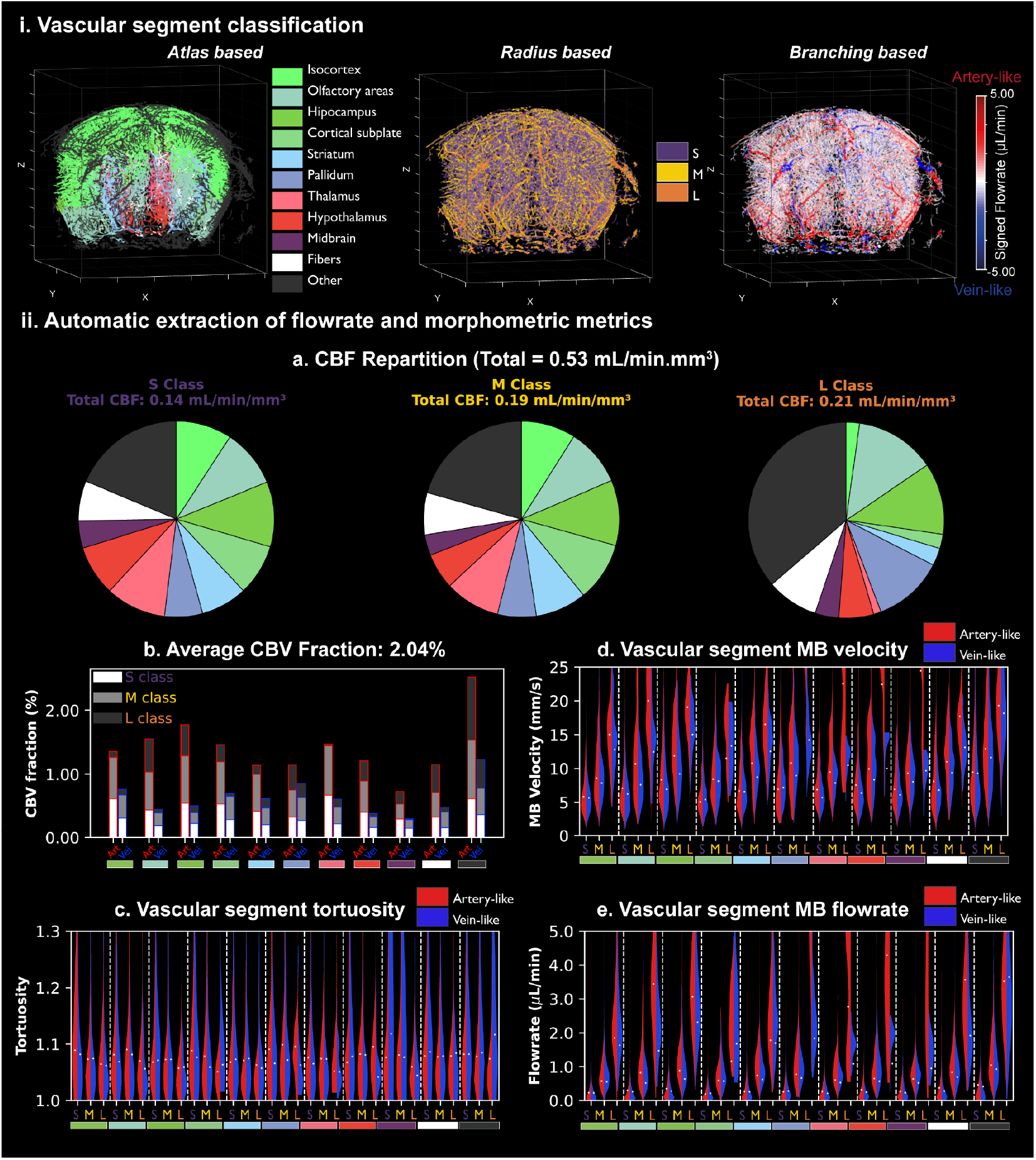
Automatic extraction of morphometric and flowrate parameters in different vascular compartments. (i) Sixty-six classes were defined using three different levels of classifications: anatomical (left), radius (middle) and branching (right). (ii) Results of the automatic flowrate and morphometric extractions. (ii.a) The global CBF repartition over the different anatomical regions is represented with pie plots, for each radius class (S-M-L). Each anatomical region is represented using the color-code already used in (i). (ii.b) The CBV fraction in the sixty-six different classes is represented in a bar plot figure. For each anatomical region, the left bar corresponds to the artery-like segments while the right bar corresponds to the vein-like segments. In each bar, the three sub-bars correspond to the radius-based classification (S-M-L), with distinct gray-scale colors. For each category, the distributions of the vascular segment tortuosity, MB velocity and flowrate are displayed (ii.c-e). For each anatomical region, three split-viol violin plots are represented, for each radius class. The left part (red) corresponds to the artery-like segments whereas the right part (blue) corresponds to the vein-like segments. The mean value of each distribution is indicated by the white dots.

## Results

### 3D RC-ULM imaging

The proposed RC-ULM approach allows for a precise localization and tracking of microbubbles, enabling the generation of detailed microbubble density and velocity volumes that are displayed in 3D and in 2D slices in figure 2. The MB density revealed a complex architecture of the vascular network in the mouse brain. Important arteries and veins were identified visually and are indicated in the Figure 2. Thanks to the high temporal resolution of the sequence that reaches 1000 volumes per second, we were able to reconstruct the arteries of the circle of Willis (CoW) and high velocity arteries such as the PCA with MB velocities reaching 10 cm/s, as displayed in the axial slices of the Figure 2.ii.a.. We also measured lower flowrates in small penetrating arterioles originating from the MCA and ACA branching territories, with MB velocities ranging between 1 and 10 mm/s (Figure 2.ii.b.). The binary vesselness mask and the corresponding skeleton are represented in supplementary Figures 1 and 2.

We also compared those results to the matrix-array MUX-based acquisition. Results shown in supplementary Figure 4 illustrate the gain in sensitivity between the two approaches on the same mouse with the same injection protocol.

### Spatial resolution and velocity profiles assessment

The spatial resolution of the RC-ULM volumes was first estimated at 20,26 µm, using the Fourier Shell Correlation and a 0.5 resolution cutoff (Figure 4.i). To illustrate the ability to measure the flowrate in small cortical vessels, we also selected one region of interest (ROI) around one arteriole and extracted the velocity profile at three different depths, chosen at three different levels of branching. The region of interest and the corresponding velocity profiles are shown in Figure 4.iii-iv. These velocity profiles demonstrated the ability to estimate the velocity in arterial subbranches, separated from less than half a wavelength. The FWHM and velocity amplitude peaks for each vessel crossed by each profile are reported in the legend. The minimum FWHM among the three examined profiles was 22.9 µm.

### Flow-directed graph analysis and Artery-like and Vein-like labeling

To illustrate and investigate our flow-directed graph analysis and labeling method, we first focused on two vascular territories of interest. By visually comparing our data with other microvascular volumes from the ex-vivo mouse brain vascular imaging literature^39–41^, we selected a seed segment just before the bifurcation node of the Medial Cerebral Artery (MCA) into three sub-branches, and another seed segment near to the confluence node of the two Galeno Veins (GV). These two seed segments were used to initiate the BFS algorithm, that revealed 353 descendants downstream of the MCA seed segment, delineating its perfused vascular territory, and 53 ancestor segments upstream to the GV seed segment, defining its drained vascular territory. Each resulting subgraph is presented in two different representations in the Figure 5.i-ii, the first one displaying the branching level of each vascular segment with respect to the seed position, and the second one showing the corresponding segment flowrate. While the flowrate decreases in the case of the MCA sub-graph, it increases in the case of the GV sub-graph, and the same tendency is also verified in the plots representing the evolution of the average flowrate by branching level, with an average flowrate gradient of -1.35 µL/min/mm in the MCA sub-graph case, and an average flowrate of +0.22 µL/min/mm in the MCA sub-graph case. Moreover, the average branching difference was found to be constantly positive in the arterial subgraph (MCA), and negative in the venous subgraph (GV), reflecting their respective diverging and merging behaviors along the vascular path. Using these combined metrics, 90% of the graph segments were labeled, with 5% of these labels automatically corrected to prevent biologically implausible vein-to-artery transitions. The remaining 10 % were discarded. The resulting artery-like and vein-like segments are represented in red and blue colors in the Figure 5.iii.

To validate our labelling method, we extracted a region of interest in the cortex where arterioles and venules can be recognized based on the axial velocity sign and verified that our branching classification results matched the axial velocity classification. A matching score of 85% between those two resulting graphs was obtained (Figure 5.iv). The comparisons in the MCA and GV regions are presented in supplementary Figure 3.

### Automatic extraction of morphometric and flowrate features by vascular compartments

#### Graph and label rendering

Labelization according to the three different criteria (branching, atlas and diameter) can be visualized directly on the 3D graph renderings (Figure 6.i) whereas morphometric and flowrate features for each vascular class are presented using pie plots, bar plots and split violin plots (Figure 6.ii).

#### Regional flow rate differences

Total CBF was measured at 0.53 mL/min/mm^3^. Although MB velocity and flowrate distributions (figure 6.ii.d-e) were most importantly correlated to vessel diameter, we still found important regional differences. Higher flow values were detected on the thalamus, hypothalamus, midbrain and olfactory areas. In these regions, flowrate in the artery-like vessels of the L class exceeded that of the corresponding vein-like vessels, which remains consistent with the presence of major feeding arteries passing through the areas from the CoW. CBF was more homogenous across brain regions for the S class (standard deviation (SD) = 2.8 µL/min/mm^3^), compared to the M (SD = 4.4 µL/min/mm^3^) and L (SD = 7.5 µL/min/mm^3^) classes (figure 6.ii.a) indicating fewer regional variations for the smaller vessels. Moreover, in those smaller vessels, flowrate and MB velocity were more balanced between arterial- and vein-like segments, leading to a local perfusion role.

#### Volumes of different vascular compartments

The mean CBV fraction was 2.04% (figure 6.ii.b). The S and M classes contributed most to this volume fraction (mean CBV fraction of 0.70 % and 0.85%, respectively) compared to the L class (0.47 %). Finally, artery-like vessels were detected more often than vein-like vessels (mean CBV fraction of 1.41% and 0.61%, respectively). Small and medium vessels also exhibited slightly higher tortuosity than larger ones (figure 6.ii.c).

## Discussion

The cerebral microvasculature is central to both neurophysiology and pathology, yet in vivo characterization remains a formidable technical challenge even in preclinical research. Ultrasound Localization Microscopy (ULM) has recently emerged as a promising microvascular imaging technique, offering the potential to non-invasively map microvascular networks with super-resolution precision including local blood velocity measurements. Recent advancements have demonstrated the feasibility of 3D ULM in preclinical models, leveraging matrix array technologies^14,16,19^ and more recently multiplexed matrix arrays^20,42^ and Row-Column (RC) arrays^28,29^ to reduce channel count requirements. While those works proved 3D ULM in vivo feasibility, including in rodent brains, 3D ULM remains challenging in practice due to the massive raw data-rate throughput from the arrays, intensive reconstruction processing and the complex analysis needed to make sense of the 3D ULM data. Building on these recent advances, we thus developed an optimized RCA-based ULM framework, compatible with 256-channel scanners, that enable high quality imaging, full microvascular graph reconstruction, flow-direction labeling, and region-specific quantification. We demonstrated high-resolution, high-sensitivity imaging of cerebral vessels down to 20 micrometers and the reconstruction of a flow-directed microvascular graph from RCA-based 3D ULM to extract quantitative morphological and hemodynamic metrics across anatomical regions in the mouse brain.

Our approach is well-adapted to image diverse range of vessel sizes (20 to 200 µm) and flow velocities (about 2 to 100 mm/s) which remain limited with other 3D ULM approaches such as multiplexed matrix arrays due to the lower volume rate.

Although classified as ‘small’, the smallest vessels in this dataset have diameters above the typical capillary range (∼5–8 μm) and thus likely correspond to precapillary arterioles and postcapillary venules rather than true capillaries.

We registered the reconstructed vessel volumes to the Allen Brain atlas using a preregistered mouse brain Doppler template. The high-contrast of the 3D RC-ULM data further allows for automatic graph reconstruction using standard tools developed for optical imaging^33^. By incorporating blood velocity direction in the vascular graph, we could orientate the reconstructed graph for each segment. We could then separate artery-like and vein-like subgraphs by assessing the branching or merging behavior along the vascular path, thanks to the estimations of the flowrate gradient and the difference between inward and outward connectivity for each node. This classification was compared to known vessels in the brain and particularly in the cortex, where the direction of vertical venules and arterioles is well known from the axial velocity direction.

Together with the classification in three diameters and in template-derived brain regions, this approach allows for automatic detailed anatomical and quantitative analysis of the microvascular compartments’ features at different scales. Namely, we could effectively quantify essential microvascular parameters such as flow velocity, vessel diameter, vessel tortuosity, CBV fraction and flow rate for those vascular compartments and regions.

We further hypothesize that adding such dynamic flow information into directed microvascular graphs will further enable researchers to introduce robust and scalable tools for brain microvascular studies beyond raw ULM data volumes.

The preclinical applications of RC-ULM extend from neurovascular research to drug discovery. It could enable high-throughput microvascular phenotyping in genetic mouse models, quantification of drug-induced vascular changes, and integration of local perfusion data with molecular profiles. RC-ULM could also provide localized microvascular assessment in acute neurovascular disorders (stroke, aneurysms), neurodegenerative diseases (Alzheimer’s, Parkinson’s), tumor angiogenesis, and cerebrovascular small vessel disease.

The translational potential of this technology for clinical investigations of microcirculation is also significant, especially in perioperative settings where a skull piece is removed during the surgery. Those applications include assessing tumoral vascular network in the brain to help surgeons better define the tumor margins and guide the resection or tools in deep and small brain regions without hitting larger vessels.

A key benefit of the RCA technology is its capacity to deliver high-quality 3D ULM using fewer channels (less than 256) as opposed to traditional matrix arrays that require specialized research scanner with more than a thousand electronic channels. This simplification not only reduces the complexity but also increases the technology’s accessibility and adaptability for broader applications. Compared to a multiplexed array, also the RCA does not require any additional electronics and can be plugged directly to an ultrafast scanner with a smaller cable. Furthermore, using a multiplexor requires to transmit and to receive the channels in different steps which leads to a strong diminution of the volume rate by a factor up to 16 when using a 1:4 multiplexor. In our own tests, RC-ULM consistently outperformed the multiplexed matrix array of similar frequency in vessel imaging quality, especially in high flowrate regions like the CoW (Supplementary figure 4).

There remain limitations to the RC-ULM approach. For one, while many steps from beamforming to prelocalization could be implemented in real time on the scanner GPUs in this study, performing 3D ULM including tracking and motion correction remains computationally intensive. To address these challenges, we are exploring the development of deep learning algorithms and online GPU-based tracking algorithms. Another limitation is the restricted “en face” field of view of the RCA due to intrinsic inability to steer the row and column beams on the same side. To overcome this limitation especially for clinical applications such as cardiac imaging, researchers have proposed the use of diverging lens^43–45^ or curved RCA arrays^46^. Finally, another limitation of the RC-ULM approach is its reduced flexibility in correcting skull-induced aberrations. While individual row and column corrections is possible along each dimension, full 2D adaptive correction as with dense matrix arrays remain impossible. In our case, we observed reduced imaging quality below the main skull sutures, it remains to be understood if this is due to wavefront distortions from said structures and if they could be further corrected to improve imaging in those areas.

Overall, RC-ULM with directed graph reconstruction demonstrates robust capability for high-resolution imaging and whole-brain microvascular quantification in mice compatible with preclinical functional ultrasound systems. This framework could enable large-scale acquisition and analysis of 3D ULM, facilitating comprehensive microvascular flow quantification for preclinical neurovascular research and complementing existing imaging techniques.

## Acknowledgements

This study was supported by the MICROVASC project funded by the European Innovation Council and SMEs Executive Agency under grant agreement No 101070917. We acknowledge the ART (Technological Research Accelerator) biomedical ultrasound program of INSERM.

## Author contributions

M.P, T.D, B.O, and M.T conceived the study. A.B, O.D, T.D developed sequence acquisitions, O.D, A.B, J.F acquired data. A.B, T.D, A.D., M.T and M.P. performed data processing. A.B, T.D, M.P, J.F, interpreted the results. T.D, A.B, M.P and J.F. wrote the first draft of the manuscript with substantial contribution from M.T and B.O

All authors edited and approved the final version of the manuscript. Underlying data have been verified by M.P and T.D

## Data sharing statement

MB volumes and microvascular graphs are available at a data repository on zenodo. Other data that support the findings of this study are available from the corresponding author upon reasonable request. Researchers wishing to obtain the raw data must contact the Office of Research Contracts at INSERM to initiate a discussion on the proposed data transfer or use.

