## Supplementary material for "In vivo microvascular flow quantification in the mouse brain using Row-Column Ultrasound Localization Microscopy and directed graph analysis"

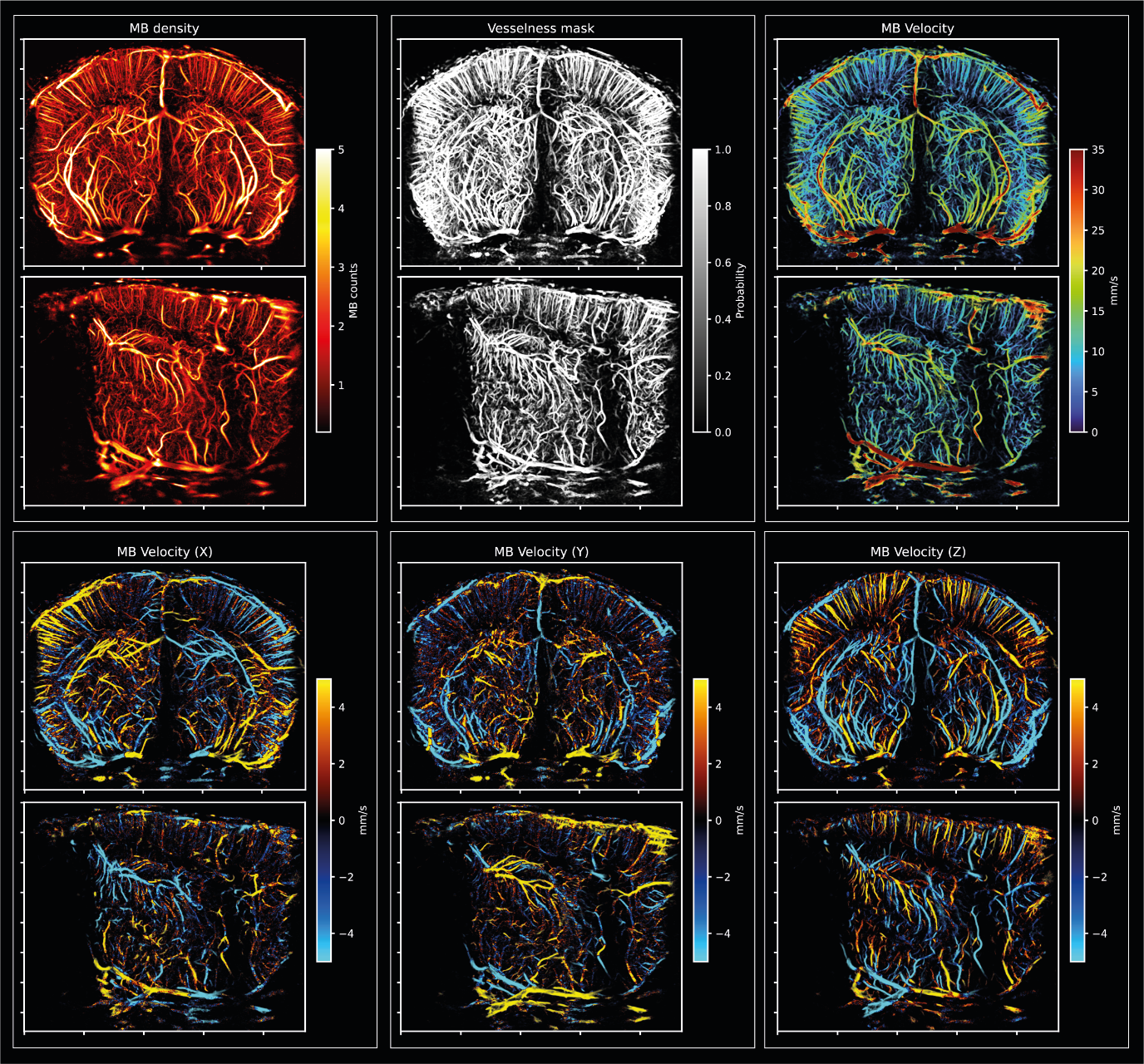
**Supplementary Figure 1:** Coronal (top) and sagittal (bottom) slices (thickness 1.5mm) of rasterized ULM data. Top left: MB density ; Top middle : Vesselness (Jerman) ; Top right : MB Velocity norm ; Bottom left : MB lateral velocity ; Bottom middle : MB antero-posterior velocity ; Bottom right : MB axial velocity. A binarized Vesselness volume was used for the 3D skeletonization, and all the ULM rasterized volume were projected on the resulting skeleton to extract the centerline value.

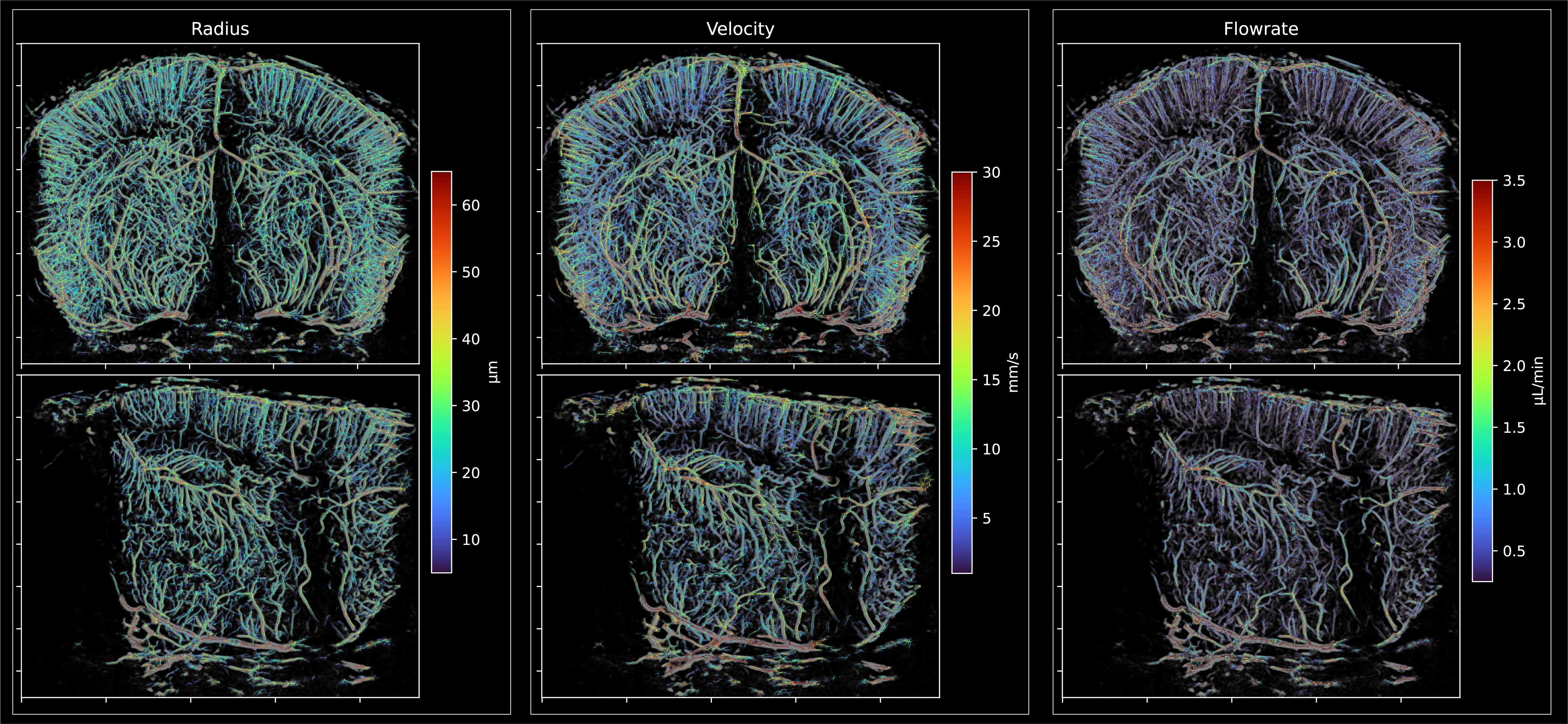
**Supplementary Figure 2:** Results of skeletonization. Different measurements are displayed on the centerline of the coronal and sagittal slices already shown in Supplementary Figure 1: Radius estimation (left), MB velocity norm (middle) and flowrate (right).

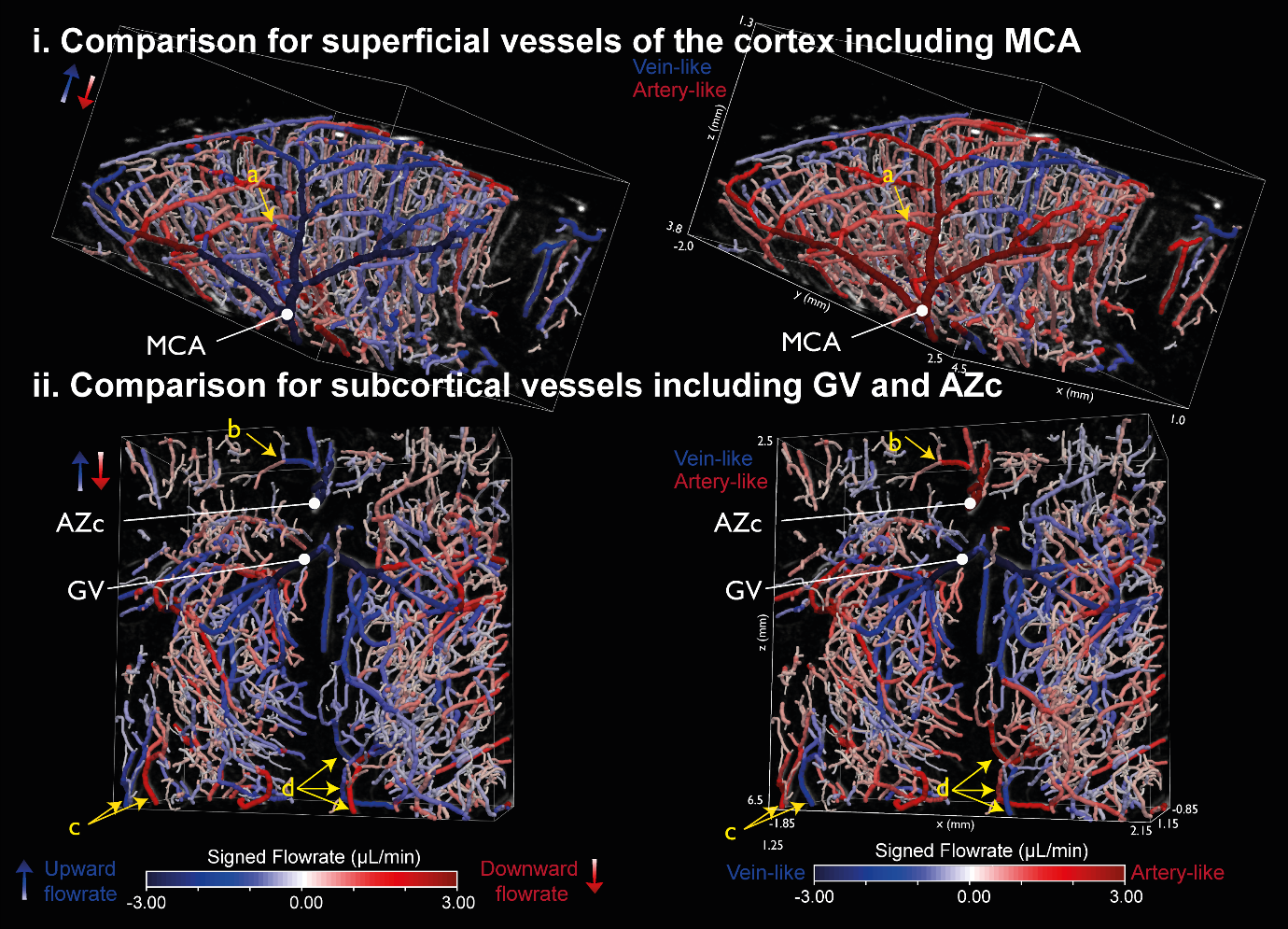

**Supplementary Figure 3:** Comparison between 3D microvascular graph renderings where the sign of the flowrate is defined by either the sign of the axial MB velocity (left), to show upward flowrate in blue and downward flowrate in red, or by the branching labelization, to show vein-like flowrate in blue and artery-like flowrate in red. **(i)** The first region contains superficial vessels of the cortex including Medial Cerebral Artery (MCA) sub-branches. These MCA sub-branches are shown in blue in the left rendering, as the MB are flowing upward in these vessels, whereas it is represented in red in the right rendering as it was well categorized as artery-like segments. Still, the penetrating arterioles in the left rendering, as the one spotted in (a) are in red. **(ii)** The second region contains sub-cortical vessels including the Galeno Vein (GV) and the Azygos Anterior Cerebral Artery (Azc). Both of them appears in blue on the left rendering as the blood is flowing upward in these vessels, however the branching classification allows to identify the GV as a vein (blue), and the Azc as an artery (red). Yellow arrows point out different artery-like vessels (red, right) that have an upward flowrate (blue, left) or vein-like vessels (blue, right) that have a downward flowrate (red, left).

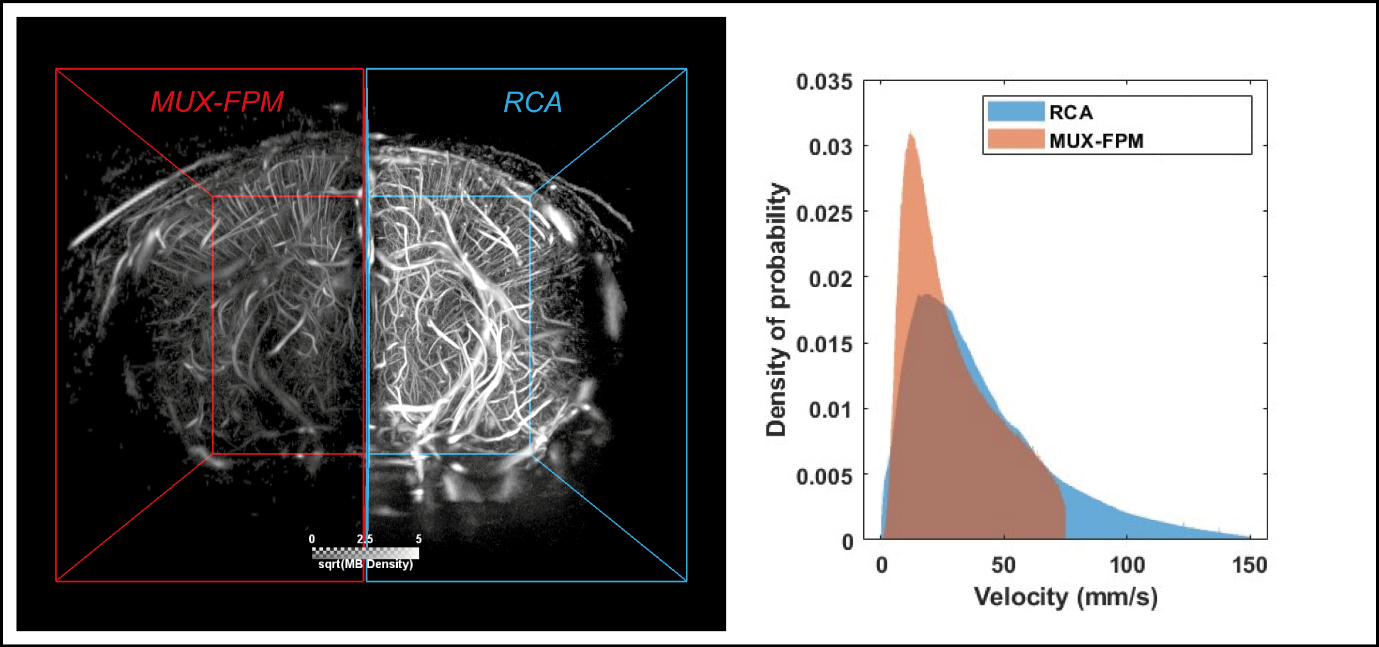

**Supplementary Figure 4:** Comparison between the RCA and the multiplexed fully populated matrix for 3D transcranial ULM in the same mouse, obtained with the same injection protocol. The RCA map show more vessels in depth, and a higher amount of MB. The velocity distributions (right panel) show higher sensitivity to fast MB for the RCA acquisition.
